# Burkholderia cepacia complex bacteremia outbreaks among non-cystic fibrosis patients in the pediatric unit of the university hospital

**DOI:** 10.1101/695247

**Authors:** Sinan Tüfekci, Birol Şafak, Özgür Kızılca, Ayşin Nalbantoğlu, Burçin Nalbantoğlu, Nedim Samancı, Nuri Kiraz

## Abstract

**Introduction:** Burkholderia cepacia complex (Bcc) leads to severe nosocomial infections particularly in the patients who have intravascular catheters and cystic fibrosis. The present study aims at investigation of Bcc outbreaks in non-cystic fibrosis patients.

**Material and Methods:** A total of 6 patients who were hospitalized at General Pediatrics Department were included in the study. Blood cultures which yielded positive signals were incubated at 5% blood sheep agar, chocolate agar and Eosin Methylene Blue agar. All fields which could be the source of the infection at the clinic were examined. Isolates confirmation with Pulsed-Field Gel Electrophoresis (PFGE) tests were performed.

**Results:** The first patient aged 14.5 years was hospitalized due to left renal agenesis, urinary tract infection and renal failure. Bcc growing was detected in blood culture which was obtained due to high fever at the 3rd day of hospitalization. New patient hospitalizations were stopped due to Bcc growing in blood cultures which was obtained due to high fever in the remaining five patients. No growing was detected in samples obtained from the clinic and the patient rooms. PFGE patterns were similar in all clinical isolates of Bcc indicating that the outbreak had originated from the same origin.

**Conclusions:** Bcc infection should always be kept in mind in nosocomial outbreaks due to multi-drug resistance and the need for hospitalization at intensive care unit. Control measures should be taken for prevention of nosocomial infections and required investigations should be done for detection of the source of the infection.

## Introduction

*Burkholderia cepacia* complex are aerobic, oxidase positive, motile, non-fermentative, spore-free gram negative bacilli which may lead to opportunistic infections. They are commonly found in soil and humid environment. *Burkholderia cepacia* complex includes at least 21 species which are phenotypically similar but genotypically different. Its identification is difficult with routine biochemical tests. Identification may be incorrect despite the presence of commercial kits. Therefore confirmation and molecular tests should be performed in reference laboratories (1). *Burkholderia cepacia* complex has recently emerged as pathogens which lead to necrotizing pneumonia and bacteriemia particularly in patients with cystic fibrosis and chronic granulomatous disease and which is intrinsically resistant to most antibiotics (2,3). However its pathogenicity is not limited with cystic fibrosis patients, it may lead to colonization and infection in respiratory tract, blood stream and urinary tract in immune compromised patients. Bcc infection was reported in intensive care units, dialysis patients, transplant patients, newborn-pediatric population and in patients with intra-venous catheters. Bcc is intrinsically resistant to aminoglycosides and polymyxine, resistant to beta-lactam antibiotics and carbapenems. Being highly contagious in hospital environment increases the importance of early diagnosis and treatment.

The present study aims at investigation of the bacteriemia outbreak in 6 patients who were hospitalized at Pediatrics Department of Tekirdag Namik Kemal University Medical School and who were detected to have *Burkholderia cepacia* complex growing in blood culture.

## Material and Methods

Six patients who were hospitalized at General Pediatrics Clinic of Tekirdag Namik Kemal University Medical School between 18 February 2019 and 03 March 2019 and who had Bcc growing in blood culture were retrospectively analyzed. Ethics committee approval was obtained prior to the study (number of decision:2019.83.06.04).

### Epidemiologic Research

Research was rapidly initiated after discussed together with Infection Control Committee as Bcc-related blood stream infection was detected in 6 patients at the same clinic. Records of the first patient were analyzed. Staff of the clinic was informed about blood culture obtaining techniques. Compliance to infection control measures was checked. Cultures were obtained from the potential infection sources, intra-venous intervention sets, antiseptics, nebulizer solutions, drugs and syringes, distilled water. Humidified sterile swabs were used. Blood was directly cultivated in 5% sheep blood agar, EMB (Eosin Methylene Blue) agar and chocolate agar and incubated at 36 ±1°C for 48 hours. Fluid samples were additionally inoculated in automatized blood culture vial and left for 5 days for incubation.

### Microbiologic Analysis

The BACTEC 9120 (Beckon Dickinson, USA) device was used for bacteria isolation. Blood culture samples which yielded positive signal were cultivated in 5% sheep blood agar and EMB (Eosin Mehylene Blue) agar and incubated at 36 ±1°C for 24-48 hours. Vitec 2 system (Bio Merieux, Marcyl Etoile, France) and conventional methods were used for identification and antibiotic susceptibility test.

Confirmation of isolates and PFGE tests were performed at Republic of Turkey Ministry of Health, General Directorate of Public Health Presidency of Microbiology Reference Laboratories and Biological Products Department. DNA extractions were done from the colonies in the medium. Afterwards DNA cut was done with Fast Digest Spel (Thermo Scientific, USA). Fingerprint was taken by using PFGE in order to investigate the clonal identity of the clinical isolates.

### Results

Of the six patients who were detected to have Bcc growing in blood culture, four were girls and two were boys. Age range was 8 months and 14.5 years. The patients did not have the history of congenital anomaly, growth and developmental retardation, chronic diarrhea, immune deficiency or cystic fibrosis. Weight and height were normal according to the age. The first patient was hospitalized due to left renal agenesis, urinary tract infection and renal failure. Bcc growing was detected in blood culture obtained due to high fever at the 3^rd^ day of hospitalization. New patient hospitalizations were stopped due to Bcc growing in blood cultures which was obtained due to high fever in the remaining five patients. The third and fourth patients required mechanic ventilation at the Intensive Care Unit as they developed respiratory failure.

The bacteriae which grew in blood cultures of the patients were identified as *Burkholderia cepacia* complex. All strains were susceptible to trimethoprim-sulphamethoxazole and resistant to ceftazidime (MIC:16 mg/L), intermediately susceptible to meropenem (MIC: 4 mg/L).

PFGE patterns were similar in all clinical isolates indicating that the outbreak was originated from a single source (Figure 1). Growing was not detected in the samples obtained from the clinic and patient rooms, so the source could not be detected. The patients who had Bcc growing in blood cultures were discharged with recovery 7-21 days after trimethoprim-sulphamethoxazole treatment. No growing was detected in control blood cultures.

**Figure 1.**
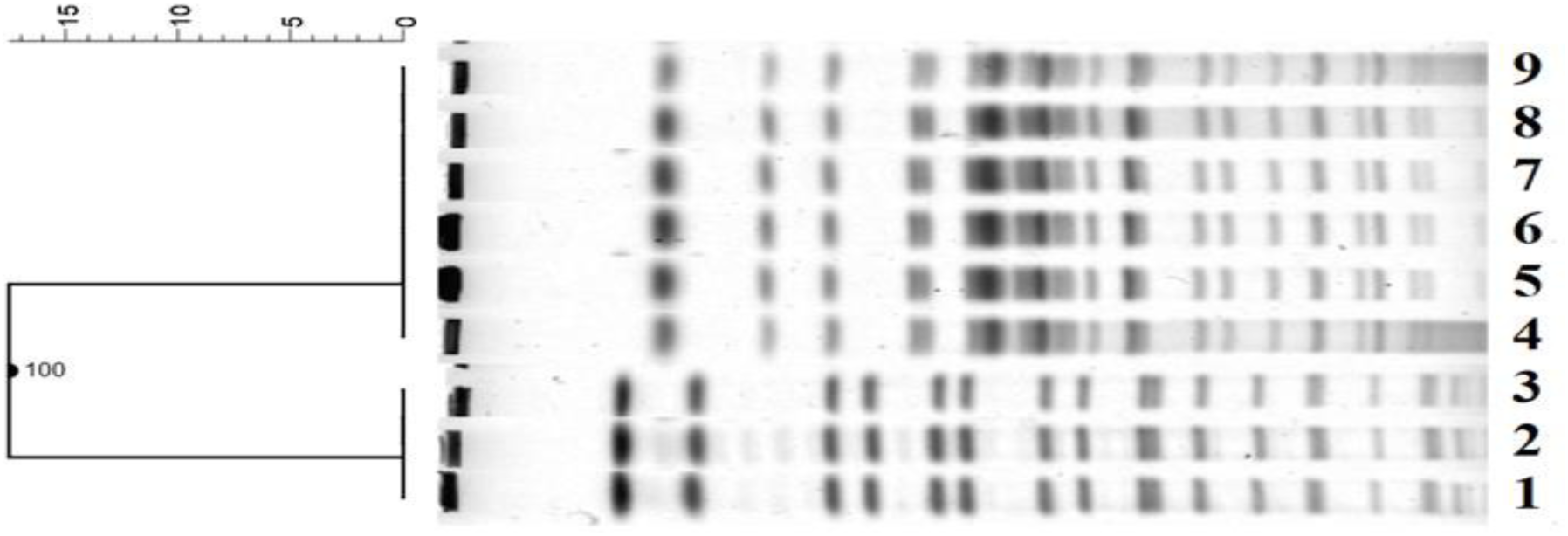
Pulsed-field gel electrophoresis patterns for 6 clinical isolates. Lane 1-3, molecular weight standarts; Lane 4-9, clinical isolates.

## Discussion

*B. cepacia* has recently been added to the non-fermentative gram negative bacteriae like *Pseudomonas aeruginosa, Acinetobacter baumannii* (4). Bcc which is found in water resources, soil, plants and nature may be detected in water resources of the hospitals, taps and sinks, various intra-venous and irrigation solutions like saline solution, nebulizer drugs, respiratory devices which uses tap water or distilled water, catheters, dialysis fluids and machines, blood gas measurement devices, termometers, ventilator heat sensors, containers which are used for enteral feeding, disinfectants and antiseptics including povidon iodine, intra-venous caffeine citrate, ultrasound gel, moisturizer, clorhexidine and benzalkonium chloride (5,6,7,8,9,10). The source of the outbreak was reported to be detected in 22 out of 30 Bcc outbreaks in non-cystic fibrosis patients (1,11,12) however the source could not be detected in the remaining eight outbreaks. The source could not be detected in the outbreak seen in Pediatrics Clinic of our hospital despite the detailed investigations done similarly with the literature.

The presence of a central venous catheter, hemodialysis-requiring renal failure, multiple bronchoscopies and recent surgeries were reported as the risk factors for Bcc bacteriemia in case-control studies. It was reported that the need for and duration of mechanic ventilator and the need for tracheostomy increased in Bcc cases compared to control group (13). Our patients did not have the history of central venous catheterization, invasive interventions, hemodialysis, surgery or bronchoscopy. Treatment took 3 weeks in the third patient who required mechanic ventilator.

Bcc may spread from one person to another directly through infected excretions and drops; indirectly through contaminated devices and equipment. Isolation of the infected patient from the others is of great importance. As the result of the outbreak at out hospital, sterilization conditions were checked, measures for isolation, hand hygiene was taken, disposable gloves and masks were used, duration of visits was decided to be shorter, health professionals’ education and environmental factors were decided to be improved.

Bcc rarely leads to an infection in healthy individuals and has a low mortality and morbidity despite having a high intrinsical resistance to many antimicrobial and antiseptic agents (14,15). Bcc may lead to life-threatening opportunistic infections like urinary tract infection, septic arthritis, peritonitis, bacteriemia, sepsis, osteomyelitis, meningitis, pulmonary abscess, pneumonia in risky patients, particularly intensive care unit patients who had an underlying disease, chronic granulomatous disease, oncologic diseases, cystic fibrosis or who are immune-compromised, who are constantly applied catheter/medical devices. It may also lead to secondary uro-genital infections due to the uro-genital interventions (6,16). Ratio of intensive care unit hospitalization was reported as 61.9% by Dizbay and 52.9% by Srinivasan. This ratio was 33% in our patients. Mortality rate was reported as 41-83% in Bcc-related infections (3,4).

While vast majority of Bcc outbreaks was originated from intensive care units, our patients were hospitalized at Pediatrics Clinic. The 10-months old female patient who was discharged after completion of cystic fibrosis and recurrent pneumonia treatment 10 days ago was suspected to be the main source however bacterial growing could not be detected.

## Conclusion

Bcc which includes nosocomial opportunistic microorganisms leads to outbreaks at intensive care units due to the natural resistance to many antibiotics. It is associated with mortality and morbidity particularly at newborn, pediatrics and adult intensive care units. Removing the main source, isolation, hand hygiene, the use of disposable gloves and masks, making short visits, education of the staff, environmental cleaning and disinfection are very important for prevention of the spread of the outbreak. In our study, outbreak could be terminated in a short time through infection control measures which were taken rapidly just after the Bcc bacteriemia which was seen in Pediatrics Clinic and of which the source could not be detected. Bcc infection should be kept in mind in the patients who are hospitalized at pediatrics clinics and intensive care units and who are resistant to treatment; and the required measures should be taken.

---

**Table 1:** Clinic features of the cases

| Cases | Gender | Age<br>(year) | First Diagnosis | Blood culture<br>positive<br>date of *B. Cepacia* | Ventilator<br>care | Duration of<br>hospital<br>stay |
| --- | --- | --- | --- | --- | --- | --- |
| 1 | female | 14, 6/12 | Left renal<br>agenesis,<br>Urinary<br>infection, | 18.02.2019 (+)<br>20.02.2019 (+)<br>22.02.2019 (+)<br>24.02.2019 (+)<br>27.02.2019 (-) | No | 12 |
| 2 | male | 14, 3/12 | Vasculitis | 20.02.2019 (+)<br>27.02.2019 (-) | No | 7 |
| 3 | male | 1, 8/12 | Pneumonia | 22.02.2019 (+)<br>02.03.2019 (+)<br>08.03.2019 (-) | Yes | 21 |
| 4 | female | 0, 8/12 | Pneumonia and<br>urosepsis | 23.02.2019 (+)<br>28.02.2019 (-) | Yes | 10 |
| 5 | female | 4, 8/12 | Pneumonia | 24.02.2019 (+)<br>28.02.2019 (-) | No | 7 |
| 6 | female | 3, 6/12 | Acute<br>bronchiolitis | 25.02.2019 (+)<br>28.02.2019 (+)<br>03.03.2019 (-) | No | 14 |

